## Supplemental Table 1 for "Hybrid seed incompatibility in Capsella is connected to chromatin condensation defects in the endosperm"

**S1 Table 1: Localization of TEs loosing DNA methylation.**

| | | Pericentromeric | Non pericentromeric | total | p-value |
|---|---|---|---|---|---|
| TEs loosing CHH | observed | 3979 | 4679 | 8658 | <0.0001 |
| | expected | 2938 | 5720 | | |
| TEs loosing CHG | observed | 5646 | 5806 | 11452 | <0.0001 |
| | expected | 3886 | 7566 | | |
| TEs loosing CG | observed | 4709 | 6181 | 10890 | <0.0001 |
| | expected | 3695 | 7195 | | |
| | Total TEs | 31517 | 61372 | 92889 | |
