## Supplemental Figure 1 for "Hybrid seed incompatibility in Capsella is connected to chromatin condensation defects in the endosperm"

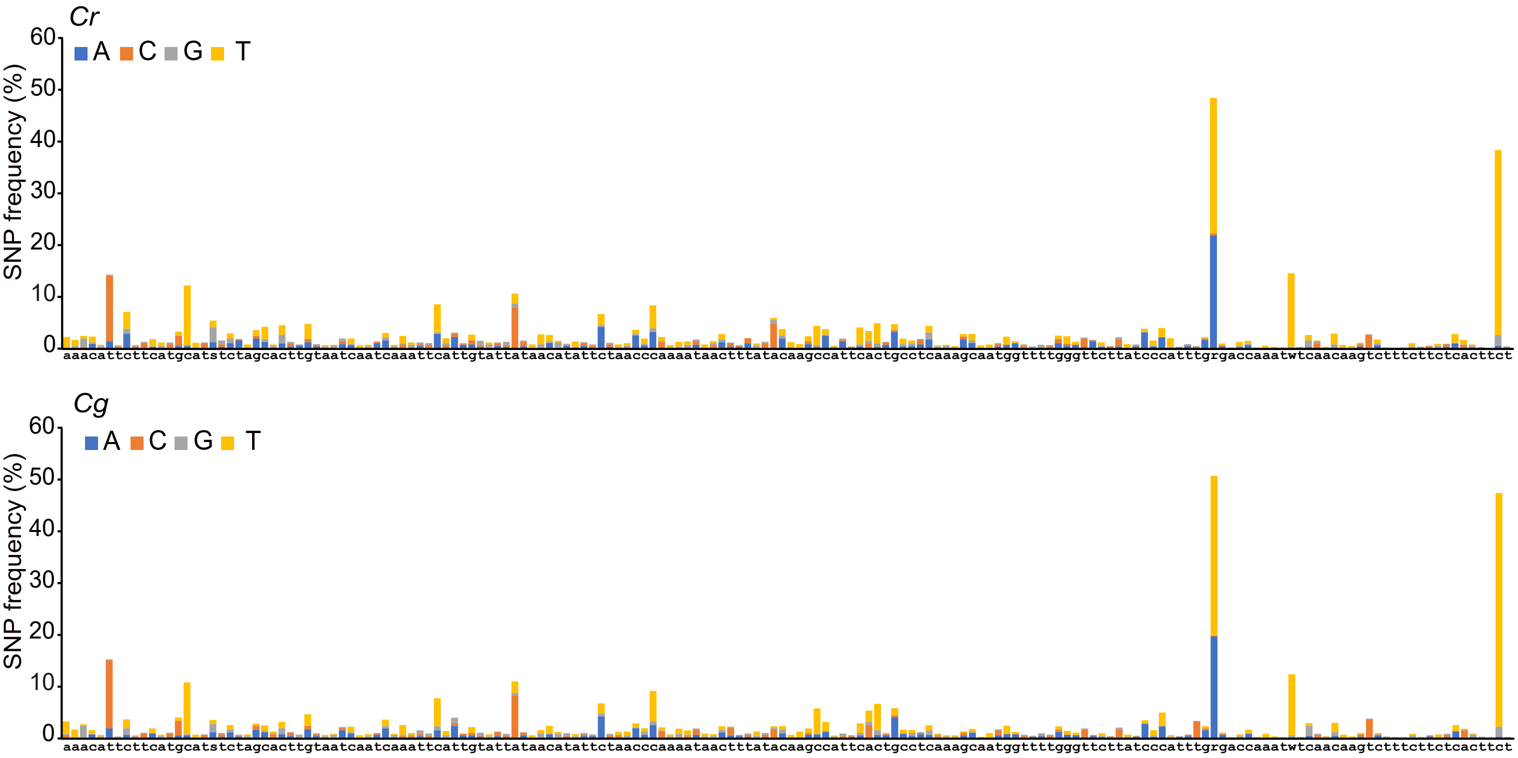


**S1 Fig: Centromeric repeats in *Cr* and *Cg* genomes are identical.**

Frequency of SNPs along centromeric repeats in *Cr* and *Cg* genomes.
