## Supplemental Figure 2 for "Hybrid seed incompatibility in Capsella is connected to chromatin condensation defects in the endosperm"

| Score | Expect | Method | Identities | Positives | Gaps |
|---|---|---|---|---|---|
| 351 bits(901) | 5e-131 | Compositional matrix adjust. | 176/178(99%) | 177/178(99%) | 0/178(0%) |

```
Cr_CENH3  1    MARTKHFATRSGPRTPADASSSQAARPSSSPATRGREGVDAAAQQPTPATSPATAKKKGA   60
Cg_CENH3       MARTKHFATRSGPRTPADASSSQAA PSSSPATRGREGVDAAAQQPTPATSPATAKKKGA
consensus 1    MARTKHFATRSGPRTPADASSSQAAVPSSSPATRGREGVDAAAQQPTPATSPATAKKKGA   60

Cr_CENH3  61   KRARFGRPQASQKKKPYRYRPGTVALREIRHYQKGTSLLIPAAAFIRQVRSITNAVAPRE   120
Cg_CENH3       KRARFGRPQASQKKKPYRYRPGTVALREIRHYQKGTSLLIPAAAFIRQVRSIT+AVAPRE
consensus 61   KRARFGRPQASQKKKPYRYRPGTVALREIRHYQKGTSLLIPAAAFIRQVRSITDAVAPRE   120

Cr_CENH3  121  VNRWTAEALVALQEAAEDFLVGLFSDSMLCAIHARRVTLMRKDFDLARRLGGKGRPW*   178
Cg_CENH3       VNRWTAEALVALQEAAEDFLVGLFSDSMLCAIHARRVTLMRKDFDLARRLGGKGRPW*
consensus 121  VNRWTAEALVALQEAAEDFLVGLFSDSMLCAIHARRVTLMRKDFDLARRLGGKGRPW*   178
```

**S2 Fig: Protein alignment of CENH3 from *Cr* and *Cg*.**

Red arrows indicate amino acid differences between proteins. Grey areas mark the sequences used as peptides for antibody production.
