## Supplemental Figure 3 for "Hybrid seed incompatibility in Capsella is connected to chromatin condensation defects in the endosperm"

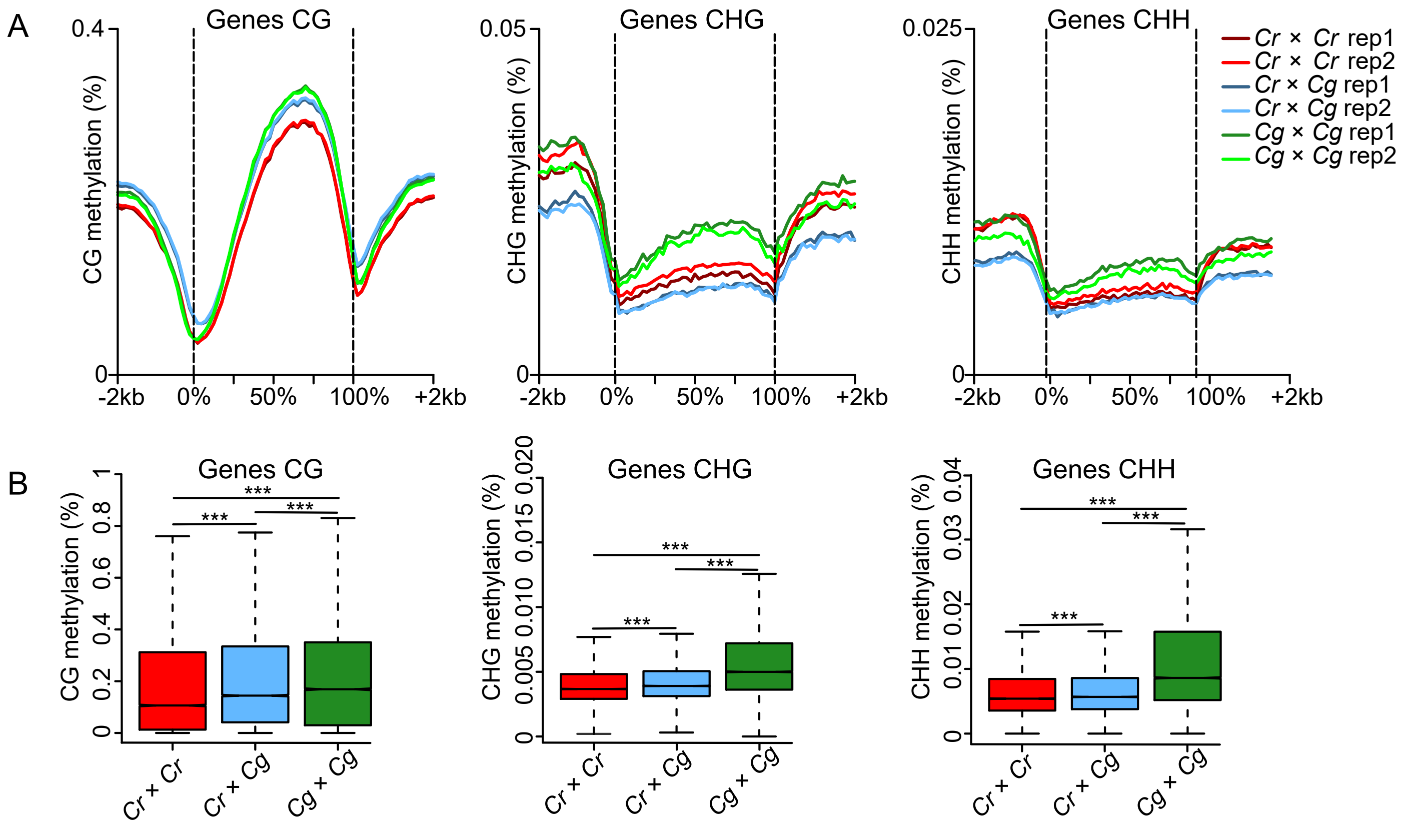


**S3 Fig: Methylation level of genes in *Cr*, *Cg* and *Cr* × *Cg* endosperm of 4 DAP seeds.**

A) Methylation level of genes in *Cr*, *Cg* and *Cr* × *Cg* endosperm of 4 DAP seeds. B) Boxplots show the methylation level of genes in *Cr*, *Cr* × *Cg* and *Cg* endosperm of 4 DAP seeds. Asterisks indicate significant differences calculated by Wilcoxon's test (*** p-value < 0.001).
