## Supplemental Figure 4 for "Hybrid seed incompatibility in Capsella is connected to chromatin condensation defects in the endosperm"

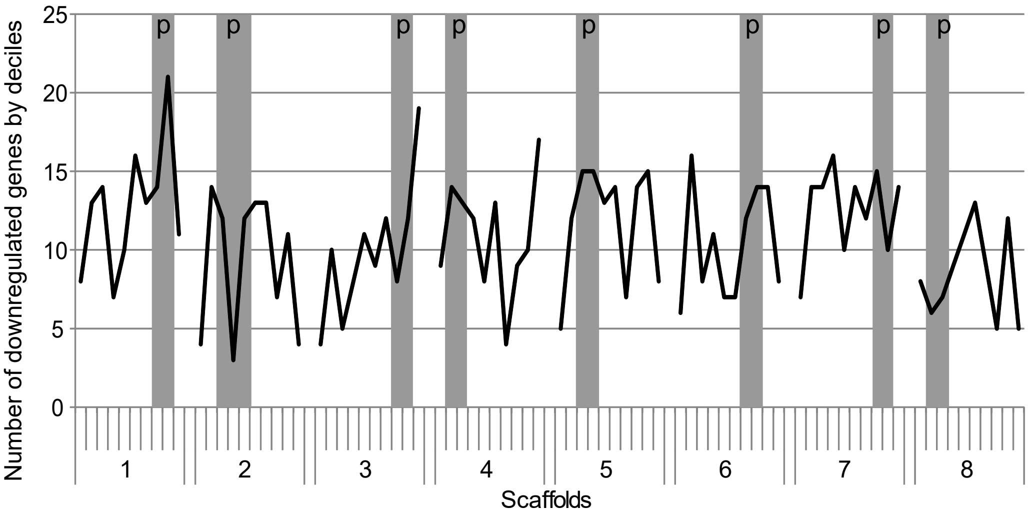


**S4 Fig: Downregulated genes in hybrid endosperm are not preferentially localized in pericentromeric regions.**

Number of genes downregulated in *Cr* × *Cg* in comparison to *Cr* per decile of genes on each scaffold. Pericentromeric regions (p) are highlighted in grey for each scaffold.
