## Supplemental Figure 5 for "Hybrid seed incompatibility in Capsella is connected to chromatin condensation defects in the endosperm"

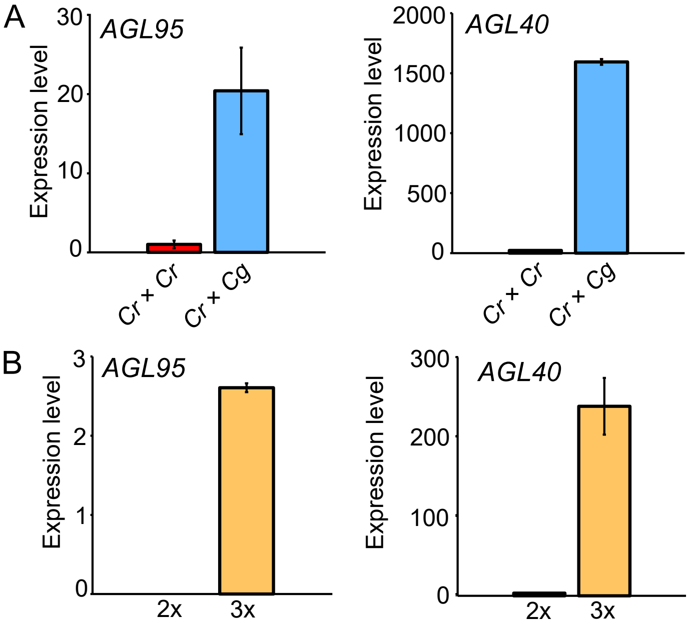


**S5 Fig: *AGL40* and *AGL95* are overexpressed in Capsella hybrid endosperm and in the endosperm of Arabidopsis triploid seeds.**

A) Expression level of *AGL95* and *AGL40* in *Cr* and *Cr* × *Cg* endosperm. *B)* Expression level of *AGL95* and *AGL40* in the endosperm of diploid and triploid seeds.
