## Supplemental Figure 6 for "Hybrid seed incompatibility in Capsella is connected to chromatin condensation defects in the endosperm"

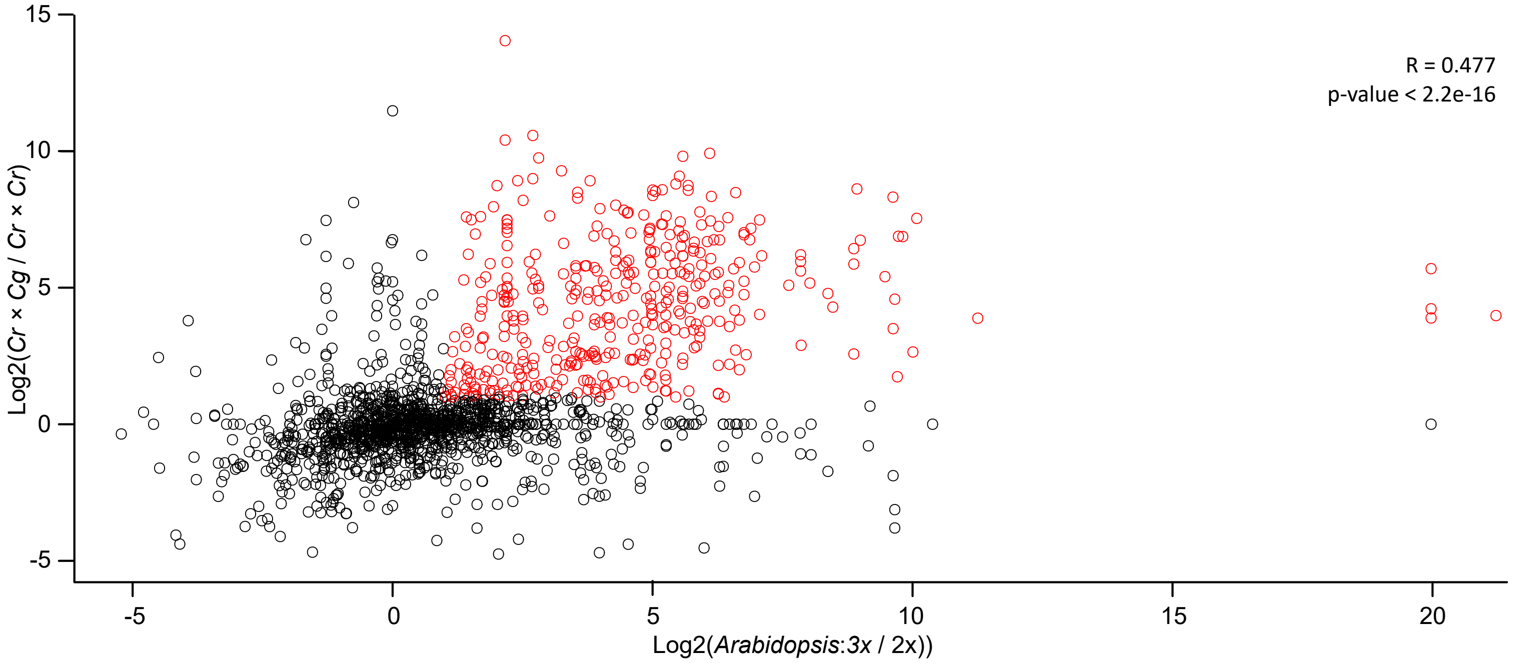


**S6 Fig: PHE1 target genes are overexpressed in Capsella hybrid endosperm and in the endosperm of Arabidopsis triploid seeds.**

Plot showing expression of PHE1 target genes in *Cr* × *Cg* compared to *Cr* × Cr seeds and in the endosperm of Arabidopsis triploid versus diploid seeds compared to diploid endosperm. Upregulated PHE1 target genes are highlighted in red.
